# Response of *Prosopis farcta* to copper and cadmium stress and potential for accumulation and translocation of these heavy metals

**DOI:** 10.1101/2020.11.02.365619

**Authors:** Mohammad Fazel Soltani Gishini, Abolfazl Azizian, Abbas Alemzadeh, Marzieh Shabani, Seifollah Amin, David Hildebrand

**Author notes:** (Corresponding authors): **Abbas Alemzadeh:**, **David Hildebrand:**.

## Abstract

Few studies have evaluated the effects of various levels of heavy metals on medicinal plants. The impact of gradually increased soil levels of copper (Cu) and cadmium (Cd) on the medicinal plant native to Southwest Asia and North Africa, *Prosopis farcta*, irrigated with metal-enriched water was determined. The exposure of plants to Cd or Cu decreased plant growth and increased Cd and Cu concentration in their shoots and roots. External Cd or Cu in the soil increased the uptake of both elements. Regression analysis showed that the weight of both shoots and roots decreased linearly with the increase of Cu and Cd contents in roots and shoots. Results showed that Cd was more toxic than Cu. The water content of shoots and roots decreased linearly with increased heavy metal levels. *P. farcta* could take up Cu and Cd in both Cu- and Cd-contaminated soils, however, it was more capable for transporting Cd from roots to shoots rather than Cu. *P. farcta* is a natural accumulator for Cu and Cd under gradually increased levels of these metals in the soil.

## Introduction

Heavy metals pollution is a major public health hazard in some locations in the world. Some metals, such as copper (Cu), are considered essential, while others, such as cadmium (Cd), are not. However, previous research shows that heavy metals, essential or non-essential, are toxic at high concentrations [1]. When plants are exposed to pollutants such as heavy metals, the plasma membrane is the first barrier to the movement of metal ions into the cytosol. Metals have been known to cause damage to plasma membranes through the binding of sulfhydryl groups of proteins and with hydroxyl groups of phospholipids [2, 3]. These metals can also replace calcium ions in cell membranes and disrupt the ionic balance of plasma membranes [4].

Plants are unique systems for monitoring cytotoxic effects induced by heavy metals stress. However, reports are sometimes conflicting or even contradictory concerning the effect of Cu on the plant defense systems against free radicals. This is due to different growth conditions, Cu concentration, types of Cu compounds, plant parts taken into consideration and different threshold levels of susceptibility to Cu stress [5, 6]. Some plants, called “accumulators”, can accumulate heavy metals in their shoots in higher concentrations than those present in the soil, without causing any damage to the plant itself [7].

*Prosopis farcta* is a medicinal plant from the Fabaceae family known as Syrian mesquite in the Middle East, and also known as Gheghe in Iran. Although little is known about optimal growth conditions of *P. farcta*, it is clear that this plant can grow very well in dry areas [8]. *P. farcta* is native to northern Africa, south western Asia, Kuwait, Turkey, Iraq and Iran [9] and is widely distributed throughout North Africa and the Middle East, especially in dry areas of most African and Asian countries. This species is widespread from India to Iran and is found in Cyprus, Turkey, Ukraine and in countries along the North African coast such as Algeria and can become naturalized in places such as Arizona [8, 10]. It is a common weed in cotton fields [11] and is also considered a host for parasitic weed like Cuscuta spp, Cistanche spp and plicosepalus acacia [12]. *P. farcta* often grows as a small shrub ranging from 30 to 80 cm tall occasionally a small tree growing up to 3 m [8]. Pods of *P. farcta* are dull black with a distinctive shape and contain 6-10 seeds [13].

It has been shown that this species is well adapted to grow under adverse environmental conditions [14]. It has been reported that *P. farcta* is a Cu-tolerant plant that grows in Cu-rich regions, such as mine areas, and can accumulate Cu in its roots and leaves, in which its chloroplasts can store approximately two times more than that of vacuoles [15]. It has also been shown that when plants are grown in contaminated soils, the chlorophyll content is increased in the plants’ leaves. It was also shown that antioxidant enzyme activities increased significantly in plants growing in contaminated soils compared to those growing in the control soil [15]. Although different parameters were investigated in *P. farcta* grown in Cu-contaminated soils [15] the effect of gradually increased levels of metals in the soil has not been thoroughly investigated.

For this reason, the overall goal of the research was to study the amplitude of growth response of *P. farcta* by quantification of fresh and dry weight after exposure to different concentrations of copper (Cu^2+^). In addition, to compare the effect of Cu with Cd, the plants were exposed to different concentrations of Cd chloride and the same parameters were measured.

## Material and Methods

### Cultivation of plants and experimental design

*P. farcta* seeds were collected from Kerman Province in the south of Iran and seed dormancy was broken by treatment with sulfuric acid [16]. Treated seeds were planted in pots filled with silty loam soil, grown under a 16/8 h day/night cycle and 25/16°C day/night temperature, and irrigated every three days with distilled water [17].

After two months, the plants were treated with different concentrations of CuSO_4_.5H_2_O and CdCl_2_, separately. CuSO_4_.5H_2_O or CdCl_2_ concentrations in solution were the following: 10, 20, 40 and 80 mg L^−1^ which means 2.54, 5.08, 10.16, and 20.32 μg mL^−1^ for Cu^2+^ and 6.13, 12.26, 24.52, and 49 μg mL^−1^ for Cd^2+^. The control plants were irrigated with distilled water. Six months after treatment, the leaves and roots were harvested separately and washed with distilled water. Treatments were arranged in a completely randomized design with five replicates.

### Measurement of plant heavy metal content

Heavy metal content was measured as previously described [18]. Leaves and roots were separated after treatment, washed and dried at 65°C for 72 h, and weighed. After drying, 1 g from each sample was placed into a porcelain crucible and heated in a furnace. The temperature was gradually elevated to 550°C in 1 h, and after 3 h the ash was dissolved in 5 mL HCl (2 N) and the total volume was adjusted to 50 mL by adding distilled water. The metal content was then analyzed by atomic absorption spectroscopy.

### Evaluation of plant’s potential for phytoremediation and translocation factor

The rate of phytoremediation was measured by bioconcentration factor (BCF) and calculated as the ratio of the metal concentration in plant (C_plant tissue_) to the soil accumulated metal concentration (C_soil_) follows [19]:

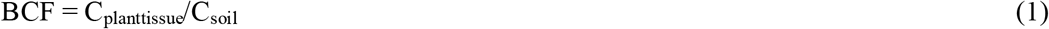

The relative bioconcentration factor (RBCF) for the two heavy metals was calculated as follows:

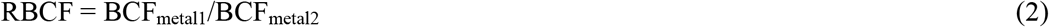

In addition, the movement of metal ions from roots to shoots is expressed in terms of the Translocation Factor (TF) [20]. TF can be calculated according to the following equation [21]:

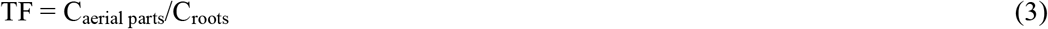

### Plant material weights and evaluation the effect of metal ions on plant growth

Whole plants were harvested and shoots and roots were separated, and then dry weight (DW) and fresh weight (FW) of roots and shoots were measured. Each part was washed separately with distilled water and weighed to determine the fresh matter, and dried in an oven at 70°C for 24 hours to measure DW.

The influence of metal ions on the growth of plants was evaluated using RFW (relative fresh weight) and RDW (relative dry weight) according to the following equations:

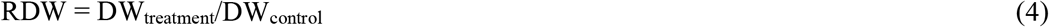

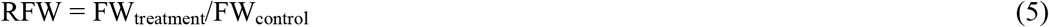

The correlation of each level of heavy metal on RDW and RFW was analyzed by linear regression.

## Results and Discussion

The Cd content of shoots and roots in the plants treated with Cd significantly increased in comparison to the control (Table 1). Interestingly, the content of Cu in the plants treated with Cd also increased in comparison to the control. This means external Cd in the soil seems to increase the uptake of Cd and Cu. Conflicting findings have been reported in different studies regarding the influence of Cd on Cu uptake by plants. In contrast to our results, one study showed that Cd treatment decreased Cu content in the shoots of corn [22], but in another study, it was reported that Cd treatment increased Cu content of corn leaves [23]. This discrepancy may be due to different growth conditions of plants. For example, in the experiment that was carried out by Azizian et al. (2013) [23], the plants were grown under drought stress conditions. For other heavy metals, it has also been reported that when the plants were exposed to a single metal, the uptake of other metals was affected [23].

**Table 1.** Contents of Cd and Cu in shoots and roots of *Prosopis fracta* exposed to different concentrations of Cd.

| **Treatment** | **Shoot** |  | **Root** |  |
| --- | --- | --- | --- | --- |
| **mg/L** | **Cd (mg/kg)** | **Cu (mg/kg)** | **Cd (mg/kg)** | **Cu (mg/kg)** |
| **0** | 1.5d | 1.4c | 1.2c | 1.6d |
| **10** | 4.8c | 4.8b | 3.2b | 5.3c |
| **20** | 5.4b | 5.1b | 3.2b | 7.2b |
| **40** | 5.8b | 4.8b | 3.9a | 7.4b |
| **80** | 8.4a | 8.0a | 4.3a | 8.8a |

Means in each column with the same letters are not significantly different at the 5% level using Duncan's multiple range test.

The Cu content of shoots and roots in Cu-treated plants significantly increased in comparison to the control (Table 2). The content of Cd in the plants treated with Cu also increased in comparison to the control. Cadmium and copper content of roots and shoots increased sharply with increasing external Cu (Table 2). These results indicate that the treatment of plants with a single metal can affect the uptake of other metals. Based on these results it can be suggested that, in this specie, there is likely the same mode of uptake of Cd and Cu. Mashhadi and Boojar [15] reported that the accumulation of Cu in the roots of *P. farcta* was 5 and 2 fold higher than levels in leaves and stems. In that study, the plants were gathered from a copper mine region in which the plants were exposed to 11,132 and 424 mg Cu/kg of soil as total and available Cu. In the present experiment, the Cu content in the shoots and roots of *P. farcta* were very similar to each other. With gradually increasing levels of Cu in the plant root media, the Cu absorbed by roots was translocated to the shoot and accumulated in the stems and leaves. Similar to these results, Snthilkumar *et al*., [24] pointed out that Cu concentration in roots and shoots of *P. juliflora* showed similar levels in plants grown in polluted soils collected from an industrial area. In contrast the Cd concentration in the plant roots was much higher than that of shoots similar to our results for *P. farcta* (Tables 1 &2).

**Table 2.** Contents of Cd and Cu in shoots and roots of *Prosopis fracta* exposed to different concentrations of Cu.

| **Treatment** | **Shoot** |  | **Root** |  |
| --- | --- | --- | --- | --- |
| **mg/L** | **Cd (mg/kg)** | **Cu (mg/kg)** | **Cd (mg/kg)** | **Cu (mg/kg)** |
| **0** | 1.6e | 1.4d | 1.0e | 1.8d |
| **10** | 2.8d | 9.8c | 1.4d | 9.4c |
| **20** | 3.1c | 9.9c | 1.8c | 11.4b |
| **40** | 3.3b | 12.3b | 2.1b | 13.0a |
| **80** | 3.7a | 14.6a | 2.4a | 14.2a |

Means in each column with the same letters are not significantly different at the 5% level using Duncan's multiple range test.

It has been shown that BCF and TF are critical in screening accumulators for phytoremediation of heavy metals [25]. Hence, these parameters were calculated for each metal to determine the potential of *P. farcta* for phytoremediation of Cu- and Cd-contaminated soils. BCF values for both heavy metals in Cd- and Cu-treated plants decreased with increasing external heavy metal levels (Table 3). This means *P. farcta* can take up these metals more efficiently when the concentration of metal is low. It has been previously reported that high concentrations of heavy metals in the soil could result in a lower BCF [26]. For example, the BCF values of about 6, 4 and 2 for *P. juliflora* grown in Cu-contaminated soil with Cu content of about 3, 6 and 7 mg/kg of soil has been reported by Sentilkumar *et al*., [24]. In addition, the BCF values show that this species can take up Cu more efficiently than Cd in both Cd- and Cu-treated plants (Table 3).

**Table 3.** Bioconcentration factor (BCF) and relative bioconcentration factor (RBCF) of Cd and Cu in Prosopis fracta exposed to different concentrations of Cd or Cu.

| Treatment<br>(mg/L) | Cd |  |  | Cu |  |  |
| --- | --- | --- | --- | --- | --- | --- |
|  | BCF-Cd | BCF-Cu | RBCF<sub>Cu/Cd</sub> | BCF-Cd | BCF-Cu | RBCF<sub>Cu/Cd</sub> |
| **0** | - | - | - | - | - | - |
| **10** | 0.4a | 0.5a | 1.3b | 0.2a | 0.9a | 4.7a |
| **20** | 0.2b | 0.3b | 1.4a | 0.1b | 0.5b | 4.4a |
| **40** | 0.12c | 0.2c | 1.3b | 0.06c | 0.3c | 4.7a |
| **80** | 0.07d | 0.1d | 1.3b | 0.03d | 0.18d | 4.7a |

Means in each column with the same letters are not significantly different at the 5% level using Duncan's multiple range test.

It has been shown that BCF is an efficient parameter that can be used to compare the potential of plants for the phytoextraction of heavy metals from polluted soil [26]. Based on our results, it can be suggested that *P. farcta* is a suitable candidate species for phytoextraction of Cu from contaminated soils. It has also been reported that BCF is more important than the content of metal ions in the plant when we want to evaluate the potential of a plant for phytoextraction [27].

In this study, for the first time a new parameter, RBCF (relative bioconcentration factor), is presented to compare the ability of *P. farcta* for taking up Cu and Cd. To use RBCF for comparisons, plants must be treated with different heavy metals and grown under the same conditions. The RBCF_metal1/metal2_ values greater than one indicate the preference of the plant to take up the first metal. In this study, RBCF_Cu/Cd_ values were higher than one, meaning this species takes up and accumulates Cu more than Cd. Our results showed that the RBCF_Cu/Cd_ values were significantly higher when plants were grown in Cu-contaminated soil (Table 3). This shows that RBCF can change when the plant is treated with different heavy metals.

In Phytoextraction, TF is also important in screening accumulator plants [26]. TF values for Cd were generally greater than Cu-TF in both Cd- and Cu-contaminated soils (Table 4). In all concentrations of heavy metals, Cd and Cu, Cd-TF values were greater than one (Table 4); this indicates that high amounts of cadmium were transported from roots to shoots. The TF value greater than one shows the translocation of the metal ions from roots to shoots [28]. It has been shown that if a species has a TF greater than one, it can be used for phytoextraction [29]. There are some reports that show hyperaccumulator plants have a TF greater than one [30, 31]. Based on these results *P. farcta* can be considered as a candidate for the phytoextraction of cadmium from contaminated soils. A similar result has been reported by Senthilkumar *et al*., [24] for the accumulation of Cd by *P. juliflora* (a species from the *Prosopis* genus) in heavy metal contaminated soils. They calculated the accumulation factor (C_plant metal_/C_soil metal_) of the Cd and Cu and concluded that *P. juliflora* is a suitable candidate for the decontamination of Cd polluted soils.

**Table 4.** Translocation Factor (TF) of Cd and Cu in *Prosopis fracta* exposed to different Concentrations of Cd and Cu.

| Treatment<br>(mg/L) | Cd |  | Cu |  |
| --- | --- | --- | --- | --- |
|  | TF-Cd | TF-Cu | TF-Cd | TF-Cu |
| **0** | 1.4c | 0.9a | 1.6b | 0.8c |
| **10** | 1.6bc | 0.9a | 2.0a | 1.0a |
| **20** | 1.8ab | 0.7b | 1.8b | 0.9bc |
| **40** | 1.5bc | 0.6b | 1.6b | 1.0ab |
| **80** | 1.9a | 0.9a | 1.0b | 1.0a |

Means in each column with the same letters are not significantly different at the 5% level using Duncan's multiple range test.

There are several proteins involved in uptake of metal ions from soil and transport to aerial parts. Some examples are carrier or channel proteins in root plasma membranes that mediate the uptake of metal ions from soil [32], such as Zip family metal transporters, [33] or another family of proteins called NRAMP (natural resistance-associated macrophage proteins) [34]. Some plant species absorb and accumulate higher amounts of heavy metals due to proteins that are specialized for this task. Based on our results, it can be suggested that there are some proteins in *P. farcta* that are involved in uptake and accumulation of Cu and Cd. These results show that this plant has an efficient mechanism(s) to transport Cd from roots to shoots, although it uses more efficient mechanism(s) to take up Cu from soils. This means there are different mechanisms in plants for uptake of heavy metals from soils and transporting them from roots to aerial parts. These results are in contrast with that of Tangahu *et al*. [31] who report that the same mechanisms exist for uptake and translocation of heavy metals in several types of plants including *Oryza sativa*, *Zea mays* and some others.

It is believed that plants accumulate heavy metals in their shoots as a defense against herbivores and pathogens through the toxicity of these metals [35]. Accumulators and hyperaccumulators are suitable candidates not only for phytoextraction but also for phytomining of precious heavy metals (such as Pt, Pd, Al and Au) [26, 36]. The use of accumulators for phytoremediation might decrease the costs of remediation [37]. Based on obtained results, *P. farcta* is a natural accumulator for Cu and Cd.

Copper is a known essential element for plant growth, but Cd is not. Copper has important roles in physiological processes, but it can be toxic to plants at high concentrations. Phytotoxic effects of Cu and Cd on plants are greater than most other heavy metals. For example, it was reported that the toxic effect of these metals on *Triticum aestivum* is greater than other metals [38]. Hence, the toxic effects of these heavy metals on *P.* farcta were determined by measuring root and shoot weights. The effects of Cu and Cd on shoot and root weights are shown in Tables 5 and 6. The weights of shoots and roots decreased under increasing concentrations of Cu and Cd. The DW decreased slower than FW with increasing heavy metal concentrations in both plants treated with Cd and Cu, which is likely due to the heavy metals comprising a larger part of plant materials in DW. In general, heavy metals act as a growth inhibitor in plants due to their adverse effects, therefore plant growth decreases when plants are grown in high concentrations of metals. Reductions of root growth with increasing external heavy metals supply levels in other plants have also been reported [39]. Although there are no previous reports on the effects of cadmium or copper on the shoot and root biomass of *P. farcta*, some reports indicate that in other plants, root and shoot weights decrease under high concentrations of Cd or Cu [40, 41].

**Table 5.** The weight of shoot and root of *Prosopis farcta* plants and their water content under Cd treatment. DW: dry weight, FW: fresh weight, DW: water content

| Cd |  |  |  |  |  |  |
| --- | --- | --- | --- | --- | --- | --- |
| Treatment (mg/L) | Shoot (g) |  |  | Root (g) |  |  |
|  | WC | DW | FW | WC | DW | FW |
| 0 | 4.1a | 3.1a | 7.3a | 3.9a | 3.2a | 7.2a |
| 10 | 3.6ab | 2.8ab | 6.4b | 3.1bc | 2.5b | 5.7b |
| 20 | 3.3bc | 2.4bc | 5.7b | 3.0bc | 2.5b | 5.5b |
| 40 | 2.8c | 2.0c | 4.9c | 2.4c | 1.9c | 4.3c |
| 80 | 2.7c | 1.9c | 4.7c | 2.4c | 1.8c | 4.2c |

Means in each column with the same letters are not significantly different at the 5% level using Duncan's multiple range test. WC: water content; DW: dry weight, FW: fresh weight.

**Table 6.**
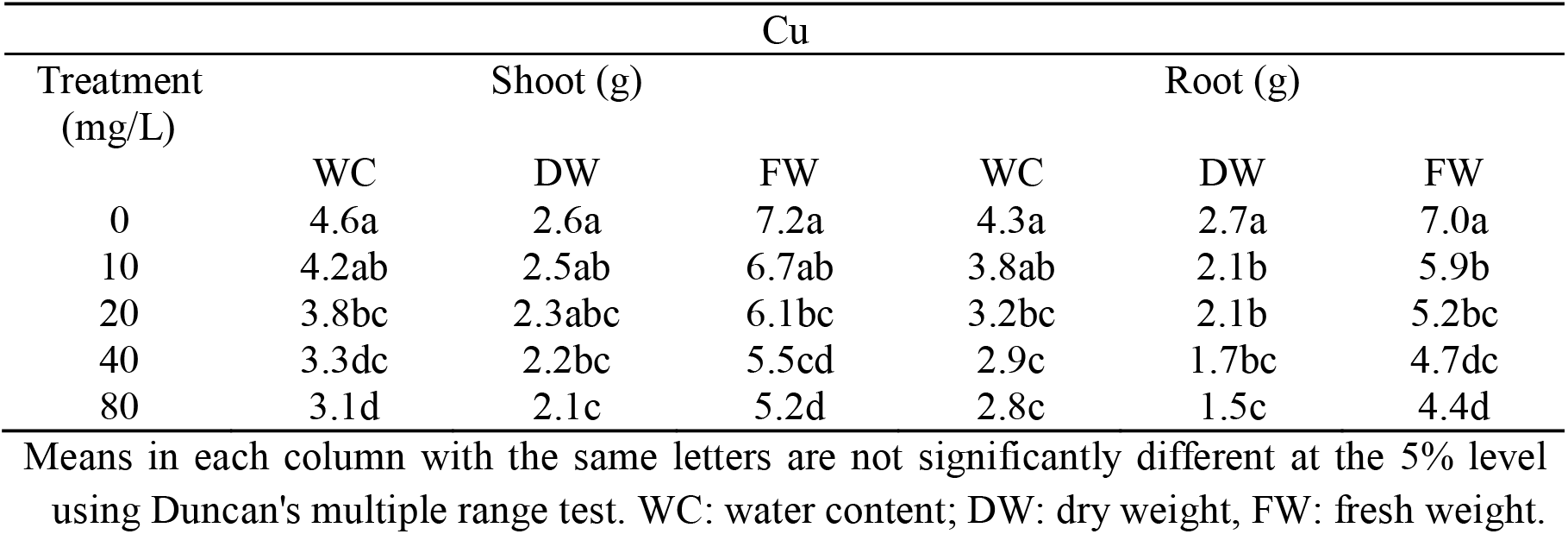
The weight of shoot and root of *Prosopis farcta* plants and their water content under Cu treatment. DW: dry weight, FW: fresh weight, DW: water content.

| Cu |  |  |  |  |  |  |
| --- | --- | --- | --- | --- | --- | --- |
| Treatment (mg/L) | Shoot (g) |  |  | Root (g) |  |  |
|  | WC | DW | FW | WC | DW | FW |
| 0 | 4.6a | 2.6a | 7.2a | 4.3a | 2.7a | 7.0a |
| 10 | 4.2ab | 2.5ab | 6.7ab | 3.8ab | 2.1b | 5.9b |
| 20 | 3.8bc | 2.3abc | 6.1bc | 3.2bc | 2.1b | 5.2bc |
| 40 | 3.3dc | 2.2bc | 5.5cd | 2.9c | 1.7bc | 4.7dc |
| 80 | 3.1d | 2.1c | 5.2d | 2.8c | 1.5c | 4.4d |

Means in each column with the same letters are not significantly different at the 5% level using Duncan's multiple range test. WC: water content; DW: dry weight, FW: fresh weight.

The water content of shoots and roots in the control plants was significantly greater than those of treated plants (Tables 5 and 6). It has been reported that Cu decreased the water content of the shoots and roots of *Vigna radiata* [42]. There are conflicting findings in different studies regarding the impact of Cd on water absorption by plants. It was observed that water absorption by kidney beans was decreased by cadmium, but that of rice and cucumber was not [43]. The decreased water absorption by corn and oats under cadmium stress has been indicated by Azizian *et al*. [44]. Another study has reported that a high concentration of Cd decreased water content of lettuce [45]. They reported that the effect of Cd seems to be due to its role in stomatal conductance, without any significant modification in net photosynthesis [45], and that it can reduce growth rate.

To determine the tolerance of plants to heavy metals, the dry or fresh weight of each treated plant was compared to control plants, and its relation to the heavy metal content of plants can be more important measures than absolute dry or fresh weights. Hence, RDW and RFW were measured to determine growth reduction, and the relation between these values and the heavy metal content of shoots and roots was analyzed by regression analysis. Regress analysis indicated that both RFW and RDW decreased linearly (p< 0.05) with the increase of Cu and Cd content in roots and shoots (Fig. 1 and 2). The results showed that both shoot and root were more sensitive to Cd than Cu (Fig. 1 and 2). In contrast to our results, it was previously observed that the root is more sensitive to Cu than other parts of rice plants [46]. This indicates that the response of plants to heavy metals differs significantly among plant species.

**Fig. 1.**
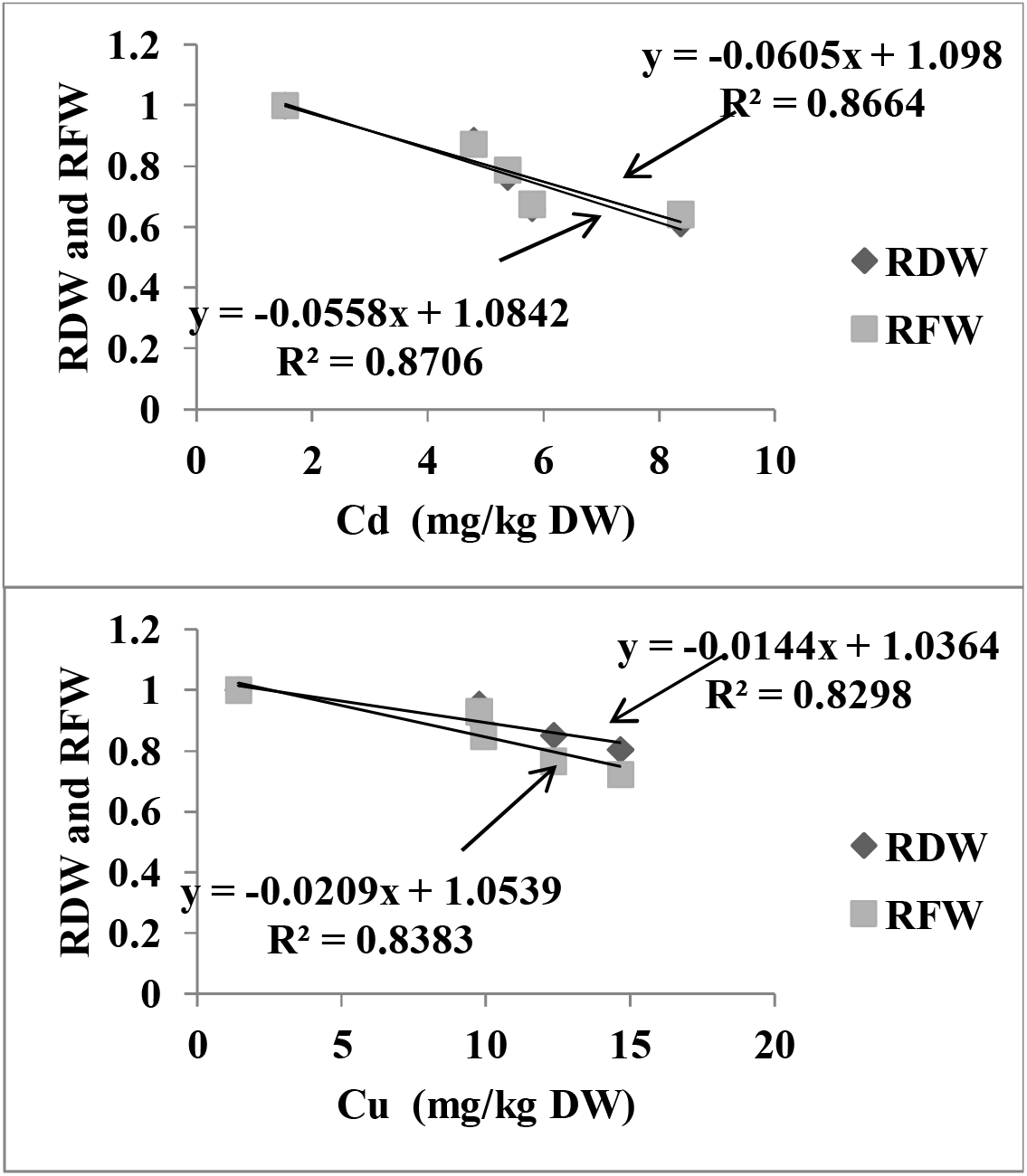
Correlation between RFW (relative fresh weight) and RDW (relative dry weight) and Cd content (up), and Cu (down) content of roots

**Fig. 2.**
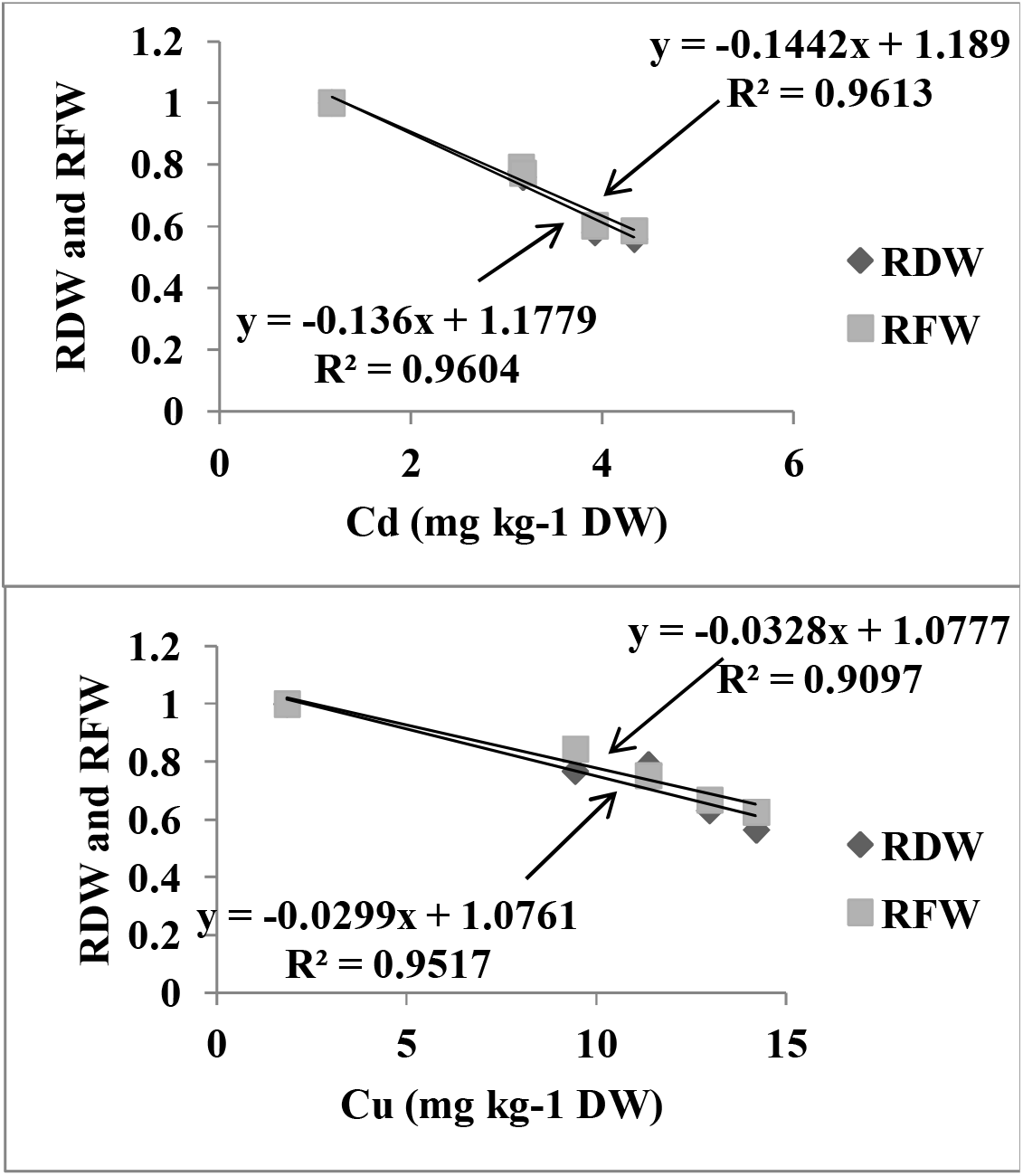
Correlation between RFW (relative fresh weight) and RDW (relative dry weight) and Cd content (up), and Cu (down) content of shoots

## Conclusions

Accumulator species are plants that take up and accumulate heavy metals in their shoots without showing toxic symptoms [7, 47]. Choosing suitable plant species for heavy metal accumulation is a critical step for successful phytoremediation of heavy metal pollutants [48]. The Brassicaceae family contains many metal accumulating species [49]. Some of these accumulators, or even hyperaccumulators, are *Thlaspi caerulescens* (Alpine pennycress) and *Alyssum bertolonii.* Alpine pennycress is one of the best-known metal hyperaccumulators [50]. *P. farcta*, which belongs to the Fabaceae family is found to also be a natural accumulator for both Cu and Cd. It has been shown previously that this species is an accumulator for Zn and Ni [51], but this is the first report to show this species is an accumulator for Cu and Cd.

TF and BCF are two important criteria for screening hyperaccumulator plants for phytoremediation of heavy metals [25, 26]. The results of the present study show that based on BCF and TF parameters, *P. farcta* can be considered a candidate for the phytoremediation of Cu- and Cd-treated soils. However, using the plant for phytoextraction of Cd and Cu should be avoided if the foliage and pods of the species are used as fodder. In addition, a new parameter, RBCF (The relative bioconcentration factor), was used in this study to evaluate the preference of a species for uptake of one heavy metal over another. *P. farcta* takes up more Cu than Cd in both Cu- and Cd contaminated soils.

In contrast to previous reports, in this study, we showed that plants probably use different mechanisms for uptake of heavy metals from soils and transporting them from roots to aerial parts. *P. farcta* transports Cd from roots to shoots, although it was more capable in taking up Cu from soils.

## List of abbreviations

Cu: Copper
Cd: Cadmium
BCF: Bioconcentration factor
RBCF: relative bioconcentration factor
WC: water content
DW: dry weight
FW: fresh weight
TF: Translocation Factor
RFW: Relative fresh weight
RDW: Relative dry weight

## Acknowledgments

This research was funded in part by grant number 82494 from Shiraz University. Special thanks to Connor Coatney, Christine Invergo and Kai Su at the University of Kentucky for editing and reviewing of this manuscript.

